# BCL6 is a context-dependent mediator of the glioblastoma response to irradiation therapy

**DOI:** 10.1101/2023.05.29.542686

**Authors:** Anna K.W. Tribe, Lifeng Peng, Paul H. Teesdale-Spittle, Melanie J. McConnell

**Author notes:** Present address: AstraZeneca, Cancer Research UK Cambridge Institute, University of Cambridge Li Ka Shing Centre, Robinson Way, Cambridge, CB2 0RE, UK. corresponding author: Email addresses. **Funding sources** This work was funded by The Cancer Society of New Zealand (Te Kāhui Matepukupuku o Aoteraroa) and Research for Life (Wellington Medical Research Foundation) (2022/338). These funding sources were not involved in the study design, collection or analysis of data, or in the writing of this report.

## Abstract

Glioblastoma is a rapidly fatal brain cancer with no cure. The resistance of glioblastoma tumours to available therapies means that more effective treatments are desperately needed. Previous research showed that the transcriptional repressor protein BCL6 is upregulated by chemo– and radiotherapy in glioblastoma and that inhibition of BCL6 enhances the effectiveness of these therapies. Therefore, BCL6 is a promising target to improve the efficacy of available treatments for glioblastoma. BCL6 is known as a transcriptional repressor in germinal centre B cells and is an oncogene in lymphoma, as well as in other cancers. However, previous research indicated that BCL6 induced by chemotherapy or irradiation in glioblastoma may not act as a transcriptional repressor. This study aimed to clarify the role of BCL6 in the response of glioblastoma to irradiation. The effect of BCL6 inhibition on the whole proteome response of glioblastoma cells to fractionated and acute irradiation treatment was investigated. Acute irradiation appeared to cause BCL6 to switch from a repressor of the DNA damage response to a promoter of stress response signalling. Rapid immunoprecipitation mass spectrometry of endogenous proteins enabled identification of proteins associated with BCL6 in untreated and irradiated glioblastoma cells. BCL6 associated with transcriptional coregulators in untreated glioblastoma and its association with the corepressor NCOR2 was validated using proximity ligation assays. However, the association of BCL6 with transcriptional regulatory proteins was lost in response to acute irradiation. This was accompanied by the irradiation-induced association of BCL6 with synaptic and plasma membrane proteins. Overall, these results reveal that the activity of BCL6 in glioblastoma therapy responses is context-dependent and may be mediated by the intensity of cellular stress.

## 1. Introduction

Glioblastoma is the most common malignant brain tumour.[1,2] It is aggressive, highly invasive and is fatal within 12 months of diagnosis in 60% of patients.[1–3] Therefore, although less common than breast, lung and colorectal cancer, the average years of life lost due to glioblastoma is much greater.[4] The standard of care for glioblastoma has not changed since the introduction of the Stupp protocol in 2005.[3,5] The Stupp protocol consists of maximal resection of the tumour followed by ionising radiation (IR) with concomitant and adjuvant treatment with the alkylating chemotherapy temozolomide (TMZ).[5] Although this regime improves survival by a few months, glioblastoma tumours display remarkable resistance to both treatments and are invariably fatal.[5]

Many factors contribute to the therapy resistance of glioblastoma, including location behind the blood brain barrier, highly effective DNA damage response mechanisms, intra– and intertumoral heterogeneity and an extremely immunosuppressive microenvironment.[6–10] It is likely that combinations of treatments targeting multiple survival mechanisms will be necessary to achieve significant survival benefits. Combining the current IR and TMZ treatments with targeted therapies that impair the ability of glioblastoma cells to survive is a potential route to improved patient outcomes. BCL6 is emerging as a promising target to improve the efficacy of glioblastoma therapies.[11–15]

BCL6 is a 79 kDa transcriptional repressor protein encoded at chromosome 3q27.[16] Homodimerisation of BCL6, mediated by the N-terminal BTB/POZ domain, is necessary for its repressive function.[17] BCL6 binds to DNA via its C-terminus zinc-finger domains and recruits corepressors to repress target gene loci.[18] These corepressors include BCOR, NCOR1 and NCOR2, which bind to the lateral groove in the BTB/POZ domain.[17,19–22] Additionally, the corepressor CtBP interacts with both the BTB/POZ domain and the middle region of BCL6, while MTA3 is recruited by the middle region.[23,24] These corepressors recruit transcriptional repression complexes containing mSIN3A and histone deacetylases (HDACs) to repress transcription.[25–28]

BCL6 is a master transcriptional regulator of the germinal centre (GC) reaction in B cells.[29–31] In GC B cells, BCL6 inhibits the DNA damage response to allow somatic hypermutation to occur without triggering cell cycle arrest and apoptosis.[29–31] Multi-layered regulation of BCL6 expression in B cells prevents the inhibition of DNA damage response pathways from becoming tumorigenic.[29–31] Constitutive expression of BCL6 in B cells leads to lymphoma and has been implicated in multiple other cancer types, including leukaemia and breast cancer.[31–37]

Several studies have implicated BCL6 in the severity and therapy resistance of glioblastoma.[11–15] BCL6 expression correlates with glioma grade.[11–15] Depletion or inhibition of BCL6 substantially decreases glioblastoma cell line viability, long-term proliferative potential and migration as well as reducing tumour growth in glioblastoma mouse models.[11,13–15] Furthermore, knockout of BCL6 in glioblastoma cell lines renders them non-viable.[11] Importantly, BCL6 inhibition enhances the effectiveness of glioblastoma therapies such as IR and TMZ both *in vitro* and *in vivo*.[11,14] Furthermore, both IR and TMZ lead to upregulated BCL6 expression.[11] Together, these observations highlight that BCL6 is vital for the survival of glioblastoma cells under unstressed conditions and plays an important role in the therapy resistance of glioblastoma.

There are indications that the role of BCL6 in glioblastoma may differ from its activity in GC B cells and lymphoma. In glioblastoma, ChIP-qPCR showed that while BCL6 bound to known target gene *TARS* and to its own exon 1, it did not bind to *TP53*, *PTEN* or *CHEK1*, which are known BCL6 targets in lymphoma.[38] When BCL6 was overexpressed in glioblastoma cells, it repressed activity of a BCL6-luciferase reporter construct as expected.[11] However, when endogenous BCL6 was upregulated by doxorubicin or IR treatment, expression of the luciferase reporter increased.[11] This suggested that endogenous BCL6 may act as a transcriptional activator in treated glioblastoma cells. Although BCL6 is known as a transcriptional repressor, BCL6-mediated transcriptional activation of *AXL* and *MED24* has been identified in glioblastoma and breast cancer respectively.[13,36] The lateral groove of the BTB/POZ domain was important for the activation of both of these genes by BCL6, suggesting that the recruitment of cofactors is involved in this activity.[13,36]

Little is known about BCL6 activity in glioblastoma or about how BCL6 promotes radioresistance. Therefore, this study investigated the effects of BCL6 inhibition on the proteome response of glioblastoma cells to both fractionated and acute IR (IR-Fr and IR-Ac). Striking differences in BCL6 activity in response to acute and fractionated treatment were investigated using rapid immunoprecipitation mass spectrometry of endogenous proteins (RIME) to identify BCL6 associated proteins.[39] The results of this study confirm that BCL6 is important in the responses of glioblastoma to IR and emphasise the context-dependency of its activity.

## 2. Results

### 2.1 BCL6 activity repressed the DNA damage response in glioblastoma cells treated with IR-Fr

LN18 glioblastoma cells were treated with daily fractions of 2 Gy ionising radiation (IR-Fr) for 5 days, to partially mimic the treatment regime administered to glioblastoma patients.[5] The effect of IR-Fr on the whole proteome of LN18 cells was investigated by label-free quantitative mass spectrometry in comparison to untreated cells. In parallel, the whole proteome of LN18 cells treated with the BCL6 small molecule inhibitor FX1 in addition to IR-Fr was compared to LN18 cells treated with IR-Fr and vehicle (DMSO).

Fig. 1A and 1B compare the proteins up– and downregulated respectively in LN18 cells in response to IR-Fr and in response to IR-Fr and FX1 (protein lists in Supplementary File 1). There were 101 proteins upregulated and 221 proteins downregulated in a BCL6-dependent fashion. These proteins only changed in abundance when BCL6 was not inhibited. A further 55 proteins were upregulated and 103 downregulated independent of BCL6 activity. The proteins only altered by IR-Fr when BCL6 was active (circled in Fig. 1A and 1B) were therefore good candidates for the BCL6-mediated response to IR-Fr and are referred to as BCL6-dependent proteins.

**Figure 1:**
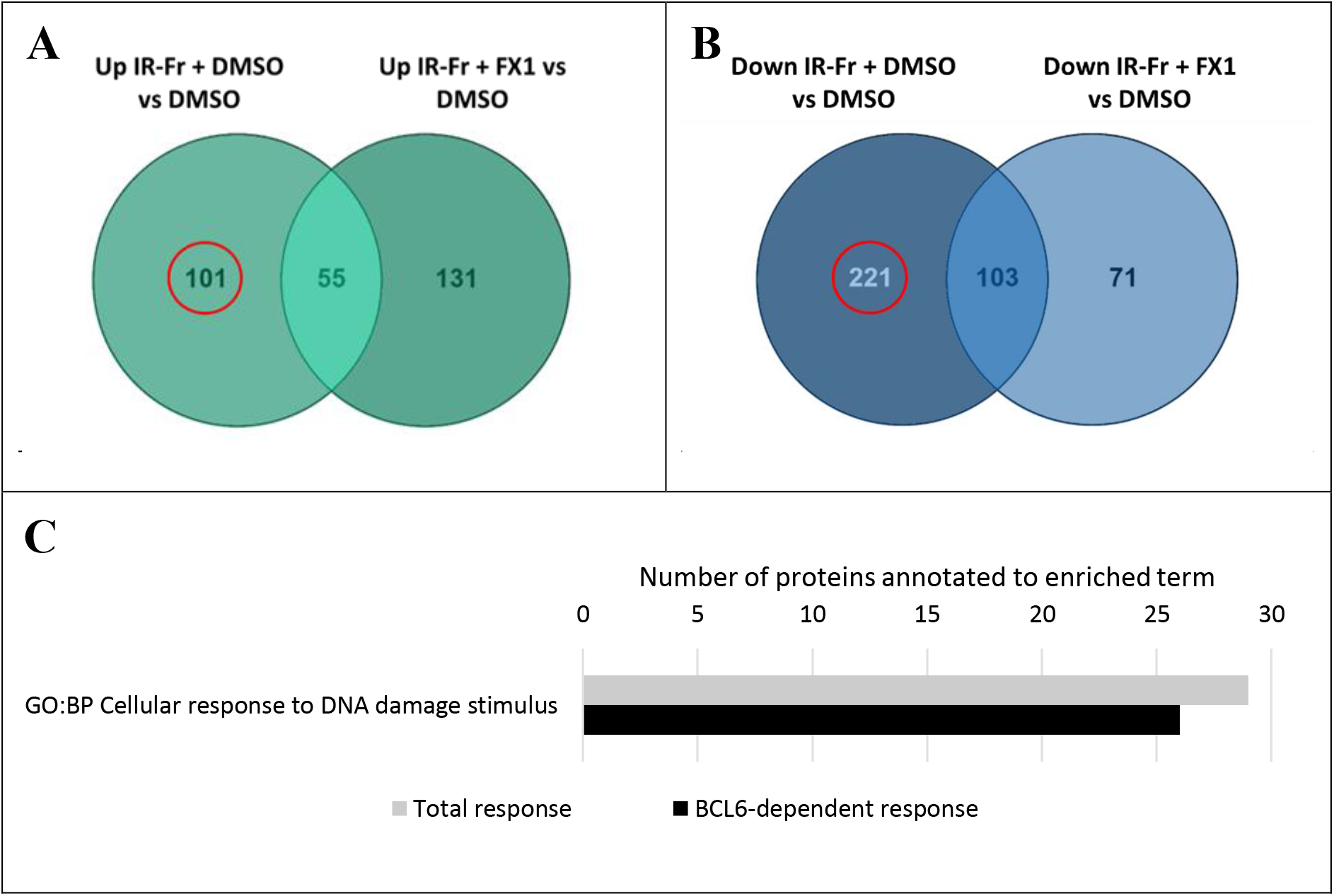
The role of BCL6 in the whole proteome response of LN18 glioblastoma cells to IR-Fr. **A)** Proteins upregulated by IR-Fr + DMSO compared to DMSO alone vs proteins upregulated by IR-Fr + FX1 compared to DMSO; **B)** Proteins downregulated by IR-Fr + DMSO compared to DMSO alone vs proteins downregulated IR-Fr + FX1 compared to DMSO. **A&B)** The BCL6-dependent proteins suggested by these comparisons are circled in red. **C)** Plot comparing the number of downregulated proteins annotated to the GO:BP term *cellular response to DNA damage stimulus* in the total LN18 whole proteome response to IR-Fr (grey) compared to in the BCL6-dependent response to IR-Fr (black).

Functional enrichment analysis was performed for the total proteome response to IR-Fr and for the subset of the response that was dependent on BCL6 (Supplementary File 2). IR-Fr treatment of LN18 cells led to downregulation of proteins involved in DNA replication and cell division and upregulation of proteins involved in translation and trafficking. These changes may have enabled LN18 cells to survive the long-term but mild therapy by slowing cell division and producing proteins needed to adapt to stress.

Strikingly, functional enrichment for the GO:BP term *cellular response to DNA damage stimulus* was not significantly enriched in the total proteome response but was enriched (p = 1.47E-2) in the analysis of the 221 BCL6-dependent proteins downregulated by IR-Fr (Supplementary File 2). Downregulation of 26 of the 29 proteins annotated to this term was dependent on BCL6 activity (Fig. 1C). These included TIMELESS and TIPIN, which are important for cell survival after DNA damage, and proteins involved in double strand break repair, such as BLM, RMI2 and NBN.[40–45] Therefore, in response to DNA damage by IR-Fr, BCL6 repressed the DNA damage response, as it does in GC B cells.[29–31]

### 2.2 BCL6 activity upregulated pathways canonically repressed by BCL6 in glioblastoma cells treated with IR-Ac

While IR-Fr is the more clinically relevant treatment, previous studies have demonstrated robust upregulation of BCL6 protein expression in glioblastoma cells 48 hours after a single 10 Gy dose of acute irradiation (IR-Ac).[11] There was no increase in BCL6 expression after IR-Fr compared to non-irradiated LN18 cells (p = 7.26E-1) (Fig. 2A and 2B), however in line with previous research, a clear trend towards increased BCL6 protein was observed in IR-Ac-treated cells (p = 8.39E-2) (Fig. 2A and 2B). Furthermore, a previous study showed that after IR-Ac treatment of glioblastoma cells, therapy-induced BCL6 did not act as a transcriptional repressor and may have acted as a transcriptional activator.[11] To investigate this apparent change in BCL6 activity, the proteome response of LN18 cells to IR-Ac with and without the BCL6 inhibitor FX1 was investigated.

**Figure 2:**
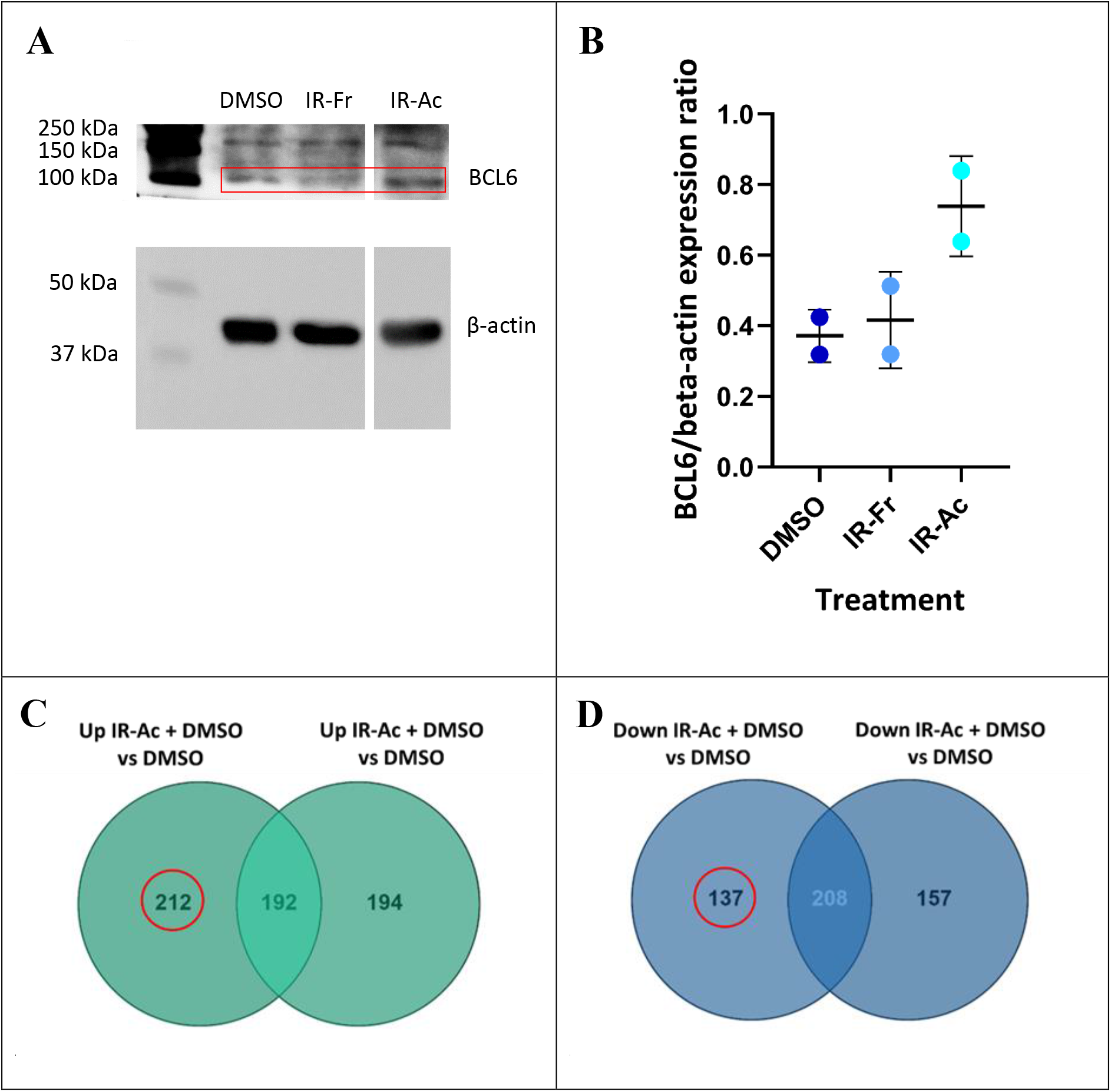
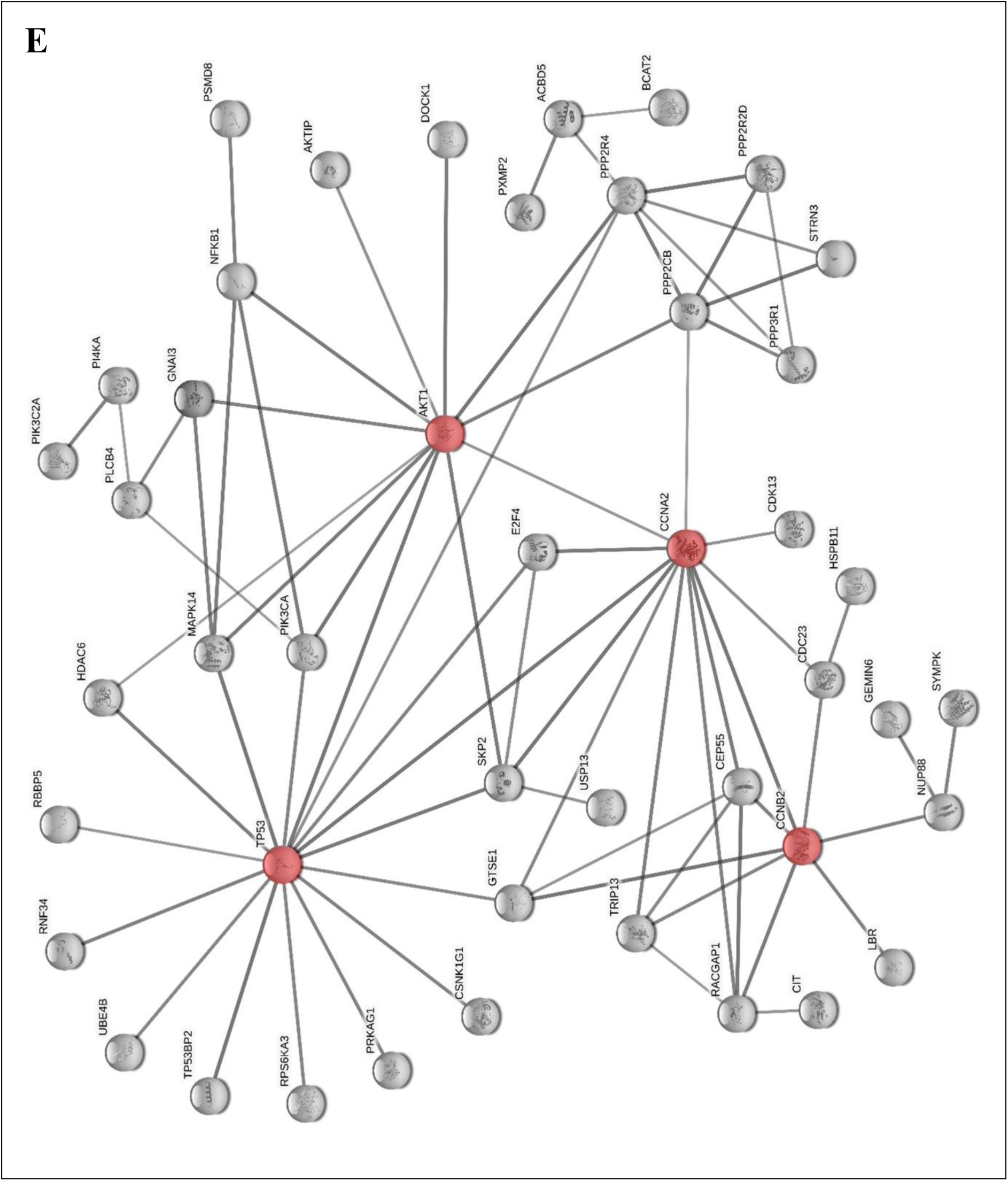
The role of BCL6 in the whole proteome response of LN18 glioblastoma cells to IR-Ac. **A)** Representative Western blot for BCL6 and β-actin in LN18 cells treated with DMSO, fractioned IR (IR-Fr) and acute IR 48 hours (IR-Ac). The western blot for BCL6 was imaged with a 5 second exposure. The membrane was stripped and re-blotted for β-actin. The western blot for β-actin was imaged with a 0.2 second exposure. **B)** Quantitative analysis of BCL6 protein relative to β-actin protein from two Western blots. Black lines show mean and error bars show standard deviation (n = 2). p values as reported in the text. **C)** Proteins upregulated by acute irradiation (IR-Ac) + DMSO compared to DMSO alone vs proteins upregulated by IR-Ac + FX1 compared to DMSO. **D)** Proteins downregulated by IR-Ac + DMSO compared to DMSO alone vs proteins downregulated IR-Ac + FX1 compared to DMSO. **C&D)** The BCL6-dependent proteins suggested by these comparisons are circled in red. **E)** STRING network of proteins dependent on BCL6 for upregulation in response to IR-Ac treatment of LN18 cells. Hub proteins highlighted in red. Edges indicate both functional and physical protein interactions and the thickness of the edges indicates confidence. Only edges with high confidence (minimum required interaction score 0.7) are shown and disconnected nodes are hidden. Interactions are sourced from textmining, experiments, databases, co-expression, neighbourhood and co-occurrence.

The whole proteome response of LN18 cells to IR-Ac indicated induction of stress responses. The proteins upregulated by IR-Ac were significantly enriched for autophagy and mitochondrial organisation terms as well as for terms related to G2/M, indicating cell cycle arrest, and to protein trafficking (Supplementary File 2). Further supporting induction of stress responses, downregulated proteins were enriched for terms related to mRNA processing and ribosome biogenesis (Supplementary File 2), suggesting a global reduction of protein turnover. Fig. 2C and 2D compare the proteins up– and downregulated respectively in LN18 cells in response to IR-Ac and in response to IR-Ac and FX1 (protein lists in Supplementary File 1). Of the upregulated proteins, 192 were upregulated regardless of BCL6 inhibition, whereas 212 proteins were dependent on BCL6 for upregulation in response to IR-Ac (Fig. 2C). Similarly, the downregulation of 208 proteins in response to IR-Ac was independent of BCL6 activity, however 137 proteins were dependent on BCL6 for downregulation in response to IR-Ac (Fig. 2D).

Unlike the BCL6-dependent response to IR-Fr, functional enrichment analysis of the BCL6-dependent response to IR-Ac did not reveal enrichment for proteins related to the DNA damage response (Supplementary File 2). Furthermore, the majority of the up– and downregulated proteins annotated to the enriched functional terms described above were not dependent on BCL6 activity.

However, STRING analysis of the proteins dependent on BCL6 for upregulation in response to IR-Ac revealed a complex network of proteins, with four clear hub proteins: p53; AKT1; CCNA2 and CCNB2 (Fig. 2E).[46] The BCL6-dependence of p53 upregulation in response to IR-Ac was surprising, as BCL6 represses p53 expression in GC B cells, lymphoma and other cancers.[32,33,35,47–49] AKT is known to be important in the radioresistance of glioblastoma, so while its upregulation in response to IR-Ac was unsurprising, its dependence on BCL6 was.[8,50–58] AKT1 was connected to various kinases and phosphatases, and to the p105 subunit of NFκB (NFκB1). BCL6 is also known to repress the NFκB pathway, so BCL6-dependent upregulation of NFκB1 was unexpected.[59] p53 also connected to kinases, as well as to E3 ubiquitin-protein ligases, transcriptional regulators and the γ subunit of the master metabolic regulator AMPK. Several of these proteins were involved in regulation of cell cycle phase transition and so were also connected to the cyclins CCNA2 and CCNB2 and other proteins involved in cell cycle control and mitosis.

This upregulation of a network of signalling proteins centred around p53, AKT1 and cell cycle regulators suggested that BCL6 was involved in the upregulation of stress response signalling in response to the DNA damage induced by IR-Ac. This is in contrast to the canonical role of BCL6 as a suppressor of the DNA damage response, which was observed in the response of LN18 cells to IR-Fr. The apparent BCL6-dependent upregulation of p53 and NFκB1, which are canonical transcriptional repression targets of BCL6, added to previous evidence that BCL6 may act as a transcriptional activator in IR-Ac treated glioblastoma cells.[11]

### 2.3 BCL6 associated with different proteins in untreated and IR-Ac treated glioblastoma cells

The striking difference in BCL6 activity in the response of LN18 glioblastoma cells to IR-Ac compared to IR-Fr merited further investigation. Based on this evidence and previous indications that BCL6 may act as a transcriptional activator in treated glioblastoma, it was hypothesised that in response to IR-Ac, BCL6 loses association with its known transcriptional corepressors, such as BCOR, NCOR1 and NCOR2, and instead associates with transcriptional coactivators.[11] This hypothesis was investigated using RIME.[39] This technique used fixation, cross-linking and immunoprecipitation to enrich endogenous BCL6 and the proteins associated with it, followed by identification of the BCL6–associated proteins using mass spectrometry.[39] The proteins associated with BCL6 in LN18 cells and in two low-passage, patient derived glioblastoma cell lines, NZG0906 and NZG1003, were investigated under untreated and IR-Ac treated conditions.[60] Three independent immunoprecipitations were carried out in each of the three cell lines, giving a total of nine replicates for untreated cells and nine replicates for IR-Ac cells. As a positive control, RIME for BCL6 was repeated in untreated Raji lymphoma cells, which endogenously express relatively high levels of BCL6.[11]

Stringent methods were used to remove non-specific and possible contaminant proteins from the RIME data. For each cell line and treatment, immunoprecipitation with a non-specific IgG control was carried out in triplicate, in parallel with the BCL6 antibody, and the associated proteins were identified. In both BCL6 and IgG control experiments, only proteins identified with high confidence (false discovery rate ≤ 0.01) were considered. First, any protein identified in the corresponding IgG immunoprecipitation was removed from the list of BCL6-associated proteins for each cell line and treatment (Supplementary File 3). Next, proteins found in fewer than 3 of the 9 untreated or IR-Ac replicates were excluded from analysis. This led to identification of proteins that were specifically and commonly associated with BCL6 in untreated and IR-Ac cells.

These data were not quantitative and simply excluded any protein found in the IgG control. For a more nuanced analysis, the proteins identified in corresponding BCL6 and IgG replicates for each cell line and treatment were quantitatively compared to find proteins that were enriched by the BCL6 antibody above the IgG signal (Supplementary File 4). If a protein had ≥ 2-fold (p ≤ 0.05) higher abundance in BCL6 replicates than IgG replicates, it was considered to be a false negative lost by the first screen. The BCL6-associated proteins identified using the quantitative method were rescued into the ‘BCL6– enriched’ set if they were enriched by the BCL6 antibody in ≥ 2/3 cell lines.

In a further control for specificity, the CRAPome database was used to identify proteins that were found in ≥ 10% of control affinity purification mass spectrometry (AP-MS) experiments.[61] These CRAPome proteins were also excluded, to ensure high confidence that the remaining proteins were truly BCL6-associated.

The EMBL-EBI tool PSICQUIC View version 1.4.11 was used to search databases conforming to the Human Proteome Organisation (HUPO) Proteomics Standard Initiative for known BCL6 protein-protein interactions (PPIs).[62] PSICQUIC View clustered the evidence for each PPI, allowing comparison of known BCL6-associated proteins with the processed RIME results. RIME for BCL6 in glioblastoma cells identified four previously recorded BCL6-associated proteins: NCOR2, TBL1XR1, FBXO11 and GOLGA2. This indicated that the technique was working as expected.

The RIME analysis identified 16 proteins commonly associated with BCL6 in both untreated and IR-Ac treated glioblastoma cells (Fig. 3A, Table 1). However, there was a clear change in BCL6 protein associations in response to IR-Ac, with 37 associations only observed in untreated glioblastoma cells and lost with IR-Ac, and an IR-Ac-induced gain of 20 new associations not seen in untreated cells (Fig. 3A, Table 1).

**Figure 3:**
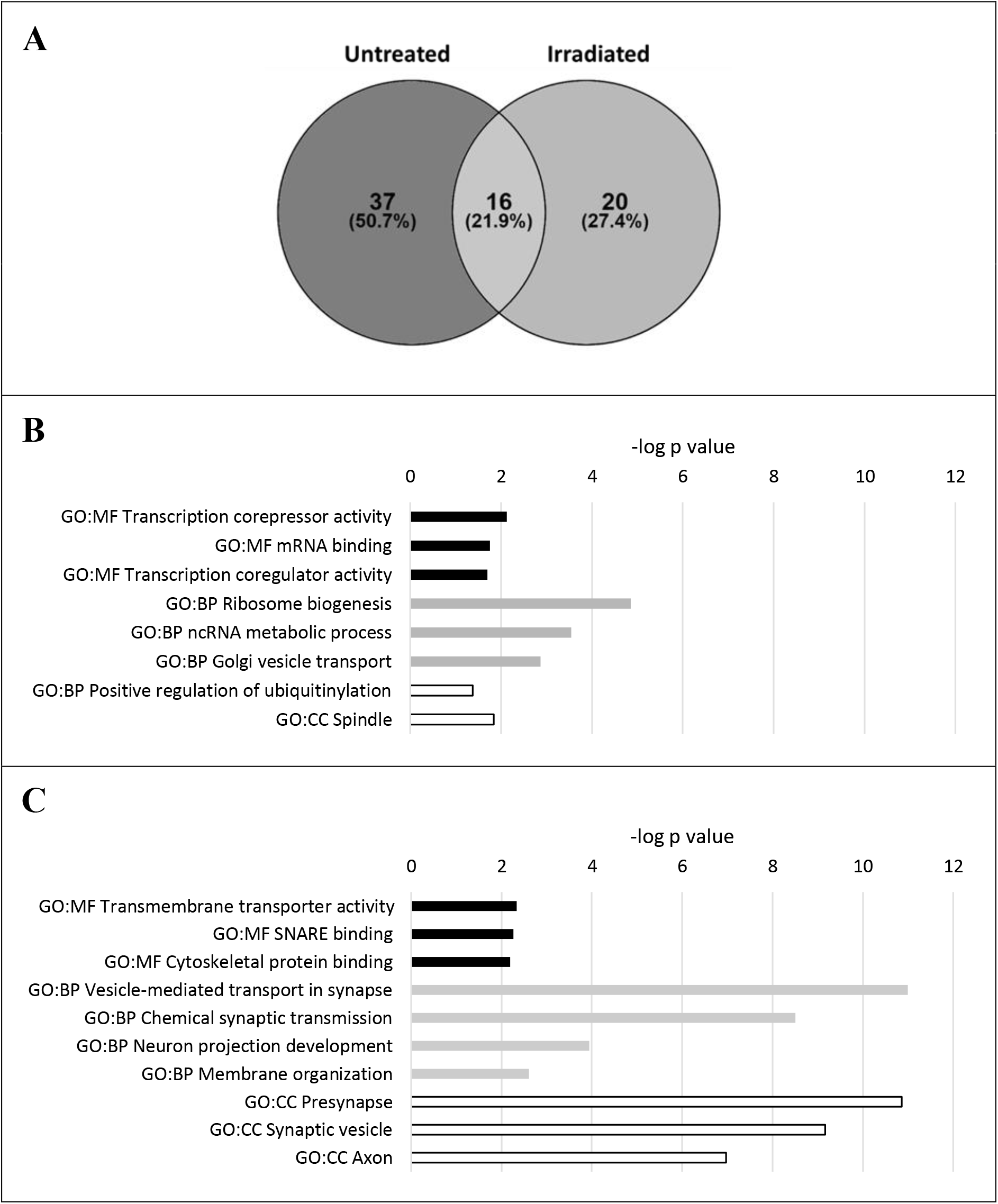
Changes to BCL6-associated proteins after IR-Ac of glioblastoma cells. **A)** Proteins commonly (3/9 replicates or 2/3 cell lines) identified as BCL6-associated in untreated and irradiated glioblastoma cells (LN18, NZG0906 and NZG1003, n=3 for each cell line and treatment). **B&C)** Up to five most significantly enriched (highest – log p value) parent GO:MF (black) GO:BP (grey) and GO:CC (white) terms from analysis of proteins that had their association with BCL6 **B)** decreased and **C)** increased by acute irradiation treatment of LN18 cells.

**Table 1:** BCL6-associated proteins in untreated and irradiated glioblastoma cells.

| BCL6-associated only in untreated cells |  | BCL6-associated in both untreated and IR-Ac treated cells |  | BCL6-associated only in IR-Ac treated cells |  |
| --- | --- | --- | --- | --- | --- |
| Protein accession | Protein name | Protein accession | Protein name | Protein accession | Protein name |
| Q01433 | AMPD2 | P04196 | HRG | P83111 | LACTB |
| Q9BZK7 | TBL1XR1 | P01024 | C3 | P02452 | COL1A1 |
| P19823 | ITIH2 | Q96LD4 | TRIM47 | Q86SE5 | RALYL |
| Q9UBF1 | MAGEC2 | P27169 | PON1 | Q06033 | ITIH3 |
| O75351 | VPS4B | Q86XK2 | FBXO11 | P61764 | STXBP1 |
| P17931 | LGALS3 | P54619 | PRKAG1 | P53680 | A2M |
| P29508 | SERPINB3 | Q13131 | PRKAA1 | Q969G5 | CAVIN3 |
| P04114 | APOB | P02461 | COL3A1 | P08123 | COL1A2 |
| P48735 | IDH2 | Q9Y618 | NCOR2 | Q03135 | CAV1 |
| Q86WA8 | LONP2 | O75146 | HIP1R | O75955 | FLOT1 |
| Q9HD26 | GOPC | P02647 | APOA1 | O94811 | TPPP |
| P51530 | DNA2 | Q08379 | GOLGA2 | P21579 | SYT1 |
| O00625 | PIR | P02458 | COL2A1 | P51674 | GPM6A |
| Q8IXK0 | PHC2 | P01599 | IGKV1-17 | P63098 | PPP3R1 |
| P42765 | ACAA2 | P02790 | HPX | Q08188 | TGM3 |
| Q99661 | KIF2C | P01023 | A2M | Q6UWE0 | LRSAM1 |
| P04040 | CAT |  |  | P16070 | CD44 |
| P12273 | PIP |  |  | Q04323 | UBXN1 |
| Q13043 | STK4 |  |  | Q6NZI2 | CAVIN1 |
| O60306 | AQR |  |  | P05026 | ATP1B1 |
| O76031 | CLPX |  |  |  |  |
| O94925 | GLS |  |  |  |  |
| P01834 | IGKC |  |  |  |  |
| Q13098 | GPS1 |  |  |  |  |
| Q6ZSZ5 | ARHGEF18 |  |  |  |  |
| Q86VS8 | HOOK3 |  |  |  |  |
| Q8IY37 | DHX37 |  |  |  |  |
| Q8IZL2 | MAML2 |  |  |  |  |
| Q8NEM2 | SHCBP1 |  |  |  |  |
| Q96KP1 | EXOC2 |  |  |  |  |
| Q99496 | RNF2 |  |  |  |  |
| Q9HC35 | EML4 |  |  |  |  |
| Q9UHA3 | RSL24D1 |  |  |  |  |
| O60701 | UGDH |  |  |  |  |
| P20930 | FLG |  |  |  |  |
| Q92504 | SLC39A7 |  |  |  |  |
| Q9HCD5 | NCOA5 |  |  |  |  |

Many of the BCL6-associated proteins identified had no previous evidence of physical interaction with BCL6. While unexpected, many of these novel findings were intriguing. For example, two subunits of master metabolic regulator AMPK, PRKAG1 and PRKAA1, were associated with BCL6 in both untreated and irradiated glioblastoma cells (Table 1). Meanwhile a variety of proteins were only associated with BCL6 in untreated glioblastoma cells, including IDH2, the pro-apoptotic, stress responsive kinase STK4 and the DNA replication and repair protein DNA2. In IR-Ac treated glioblastoma cells, BCL6 was associated with four components of caveolae, CAV1, CAVIN1, CAVIN3 and FLOT1, and with negative regulator of NFκB signalling, UBNX1. These intriguing associations and others merit further investigation in future studies as they may yield insight into BCL6 activity in untreated and IR-Ac treated glioblastoma.

Quantitative comparisons were made for the proteins associated with BCL6 in the three untreated and IR-Ac treated glioblastoma cell lines (Supplementary File 5). To investigate the amount of the proteins associated with BCL6, protein abundance was normalised to the abundance of BCL6 for each sample. After removal of non-specific and potential contaminant proteins, only the LN18 cell line had sufficient numbers of proteins with IR-Ac-altered BCL6 association (≥2-fold change, p ≤ 0.05) for informative functional enrichment analysis (Fig. 3B&C, Supplementary File 5).

Proteins that decreased association with BCL6 after IR-Ac were enriched for several gene ontology terms (Fig. 3B). Most striking was the functional enrichment for the GO:MF terms *transcriptional corepressor activity* (p = 7.50E-3) and *transcriptional coregulator activity* (p = 2.00E-2). The proteins annotated to these terms included known BCL6 binding partner NCOR2, component of NCOR complexes TBL1XR1, and other corepressors RCOR1, DRAP1 and NCOA5.[19,63,64] The *transcriptional coregulator activity* term also contained transcriptional coactivators MAML2 and PIR.[65–67]

Consistent with our hypothesis, in response to IR-Ac BCL6 decreased association with transcriptional co-regulator proteins, including known BCL6 corepressor NCOR2. However, rather than gaining associations with transcriptional coactivator proteins as hypothesised, proteins that increased association with BCL6 in response to IR-Ac were strongly enriched for GO:BP and GO:CC terms relating to synaptic activity. These terms included *vesicle-mediated transport in synapse* (p = 1.01E-11), *chemical synaptic transmission* (p = 3.11E-9), *presynapse* (p = 1.38EE-11) and *synaptic vesicle* (p = 6.82E-10), along with other terms related to neuron morphogenesis. While unexpected, this was consistent with the dramatic IR-Ac-induced changes to BCL6 activity suggested by the whole proteome analysis.

### 2.4 BCL6 was associated with some known corepressors in glioblastoma cells

The IR-Ac-induced loss of BCL6 association with transcriptional coregulator proteins in LN18 cells was examined in greater detail. The known BCL6 associated proteins HDAC1 and HDAC2 are commonly found in the negative control AP-MS datasets submitted to the CRAPome database, so they had been excluded from the initial analysis.[61] Interrogation of the full RIME data revealed that known BCL6 binding partners NCOR2, HDAC1 and HDAC2 were identified as BCL6-associated proteins in several of the untreated glioblastoma samples (Table 2). These three binding partners were also identified as BCL6-associated proteins in a handful of irradiated glioblastoma samples.

**Table 2:**
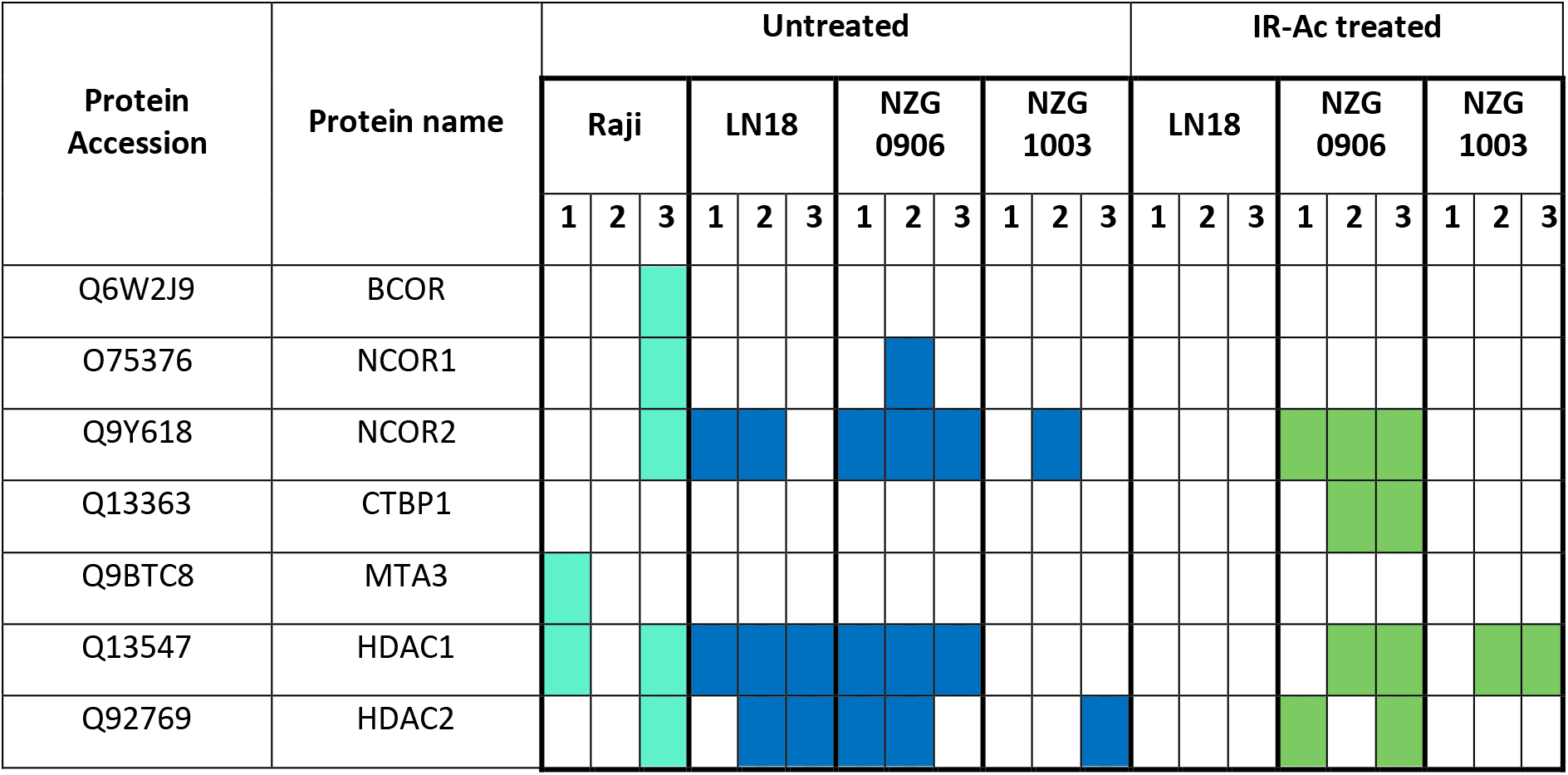
Identification of known BCL6 binding partners in RIME samples.

Other known BCL6 binding partners were identified infrequently or not at all. It is possible that BCL6 does associate with other known binding partners in glioblastoma cells but that these proteins fell below the threshold of detection. This possibility was supported by results from RIME in Raji lymphoma cells, which express a relatively high level of BCL6 protein.[11] As BCL6 has the canonical transcriptional repressor role in lymphoma, it was expected that RIME would detect BCL6 associations with known BCL6 corepressors.[31] Notably, the major BCL6 corepressors BCOR, NCOR1 and NCOR2 were only identified as BCL6-associated proteins in one of the three positive control Raji lymphoma samples (Table 2). This indicated that the signal for BCL6 in the RIME assay is at or near the limit of detection. Nevertheless, these RIME results provide the first evidence that BCL6 associates with its known binding partner NCOR2 and with HDAC1 and HDAC2 in glioblastoma cells.

Samples in which each protein was identified highlighted in turquoise for positive control Raji lymphoma samples, in blue for untreated glioblastoma samples and in green for IR-Ac treated glioblastoma samples.

### 2.5 BCL6 lost its association with transcriptional regulators in response to IR-Ac of glioblastoma cells

While BCL6 associated with known BCL6 corepressor NCOR2 in both untreated and irradiated glioblastoma samples, quantitative analysis revealed a notable decrease in the association of NCOR2 with BCL6 after IR-Ac treatment of LN18 and NZG0906 cells (Fig. 4). Although only statistically significant (p ≤ 0.05) in LN18 cells, this decrease in BCL6-NCOR2 association was most distinct in NZG0906 cells, in which the number of peptide spectral matches (PSMs) detected for each protein conferred confidence that NCOR2 was not fluctuating around the threshold of detection, as was possible in the LN18s (Table 3).

**Figure 4:**
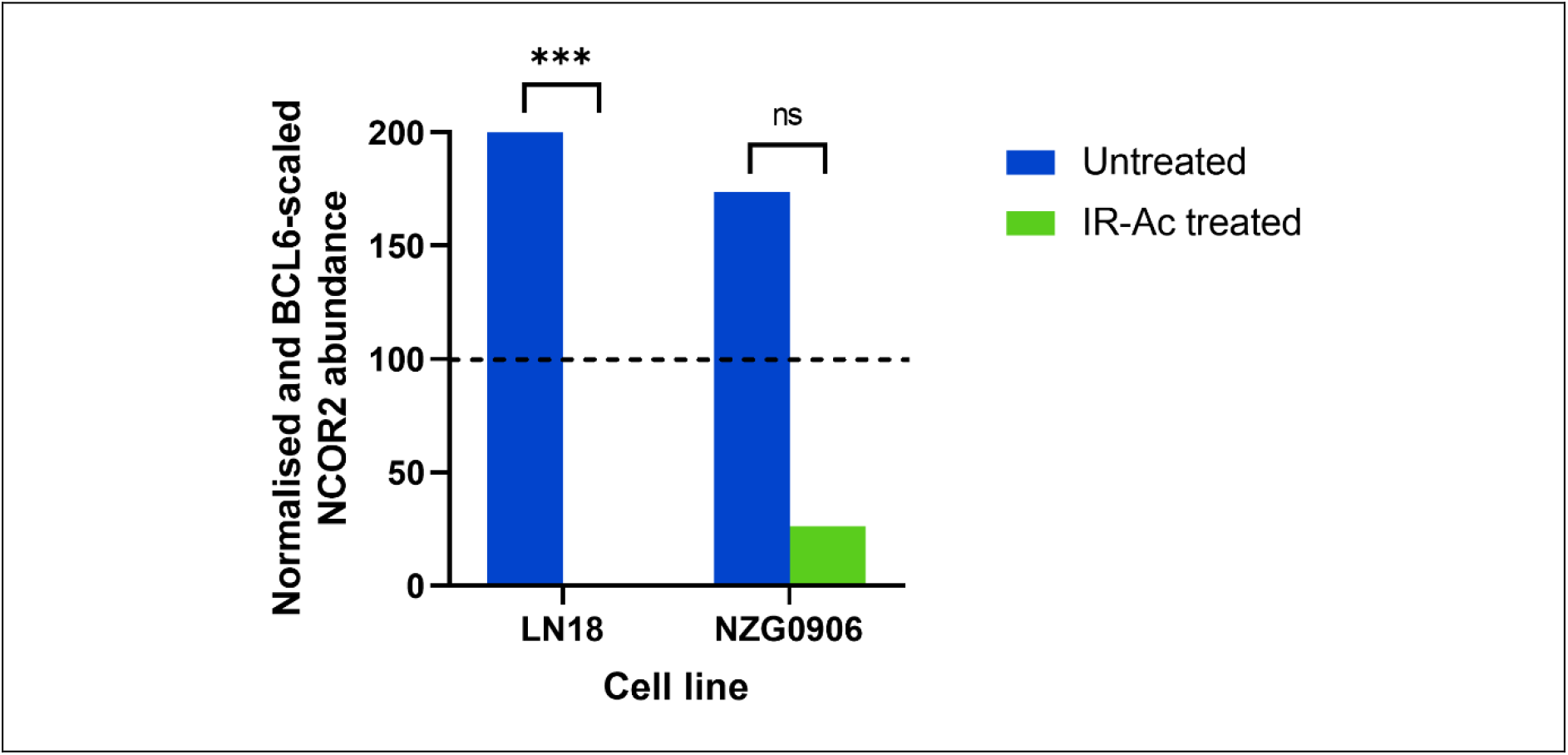
Loss of NCOR2 association with BCL6. Abundance values reflect the median of all possible pairwise ratios of peptide peak intensities between replicates, with normalisation to BCL6 abundance (dotted line) and scaling so that the average abundance of all the samples was 100. *** = p ≤ 0.01, ns = p ≥ 0.05 (non-significant). Processing settings excluded the NCOR2 PSMs identified in NZG1003 samples from quantification.

**Table 3:** Peptide spectral matches to BCL6 and NCOR2 across RIME samples.

| Cell line | Treatment | Replicate | PSMs |  |
| --- | --- | --- | --- | --- |
|  |  |  | BCL6 | NCOR2 |
| LN18 | Untreated | 1 | 2 | 1 |
|  |  | 2 | 1 | 3 |
|  |  | 3 | 1 | - |
|  | IR-Ac | 1 | 2 | - |
|  |  | 2 | 1 | - |
|  |  | 3 | 1 | - |
| NZG0906 | Untreated | 1 | 5 | 2 |
|  |  | 2 | 5 | 6 |
|  |  | 3 | 3 | 3 |
|  | IR-Ac | 1 | 13 | 5 |
|  |  | 2 | 8 | 4 |
|  |  | 3 | 13 | 2 |
| NZG1003 | Untreated | 1 | 1 | - |
|  |  | 2 | 7 | 1 |
|  |  | 3 | 2 | - |
|  | IR-Ac | 1 | 10 | - |
|  |  | 2 | 8 | - |
|  |  | 3 | 10 | 1 |

Further interrogation of the RIME data revealed the association of BCL6 with other transcriptional regulators. Most were identified from a low number of PSMs in only some RIME samples, again indicating that BCL6-associated proteins were close to the threshold of detection by RIME. BCL6 commonly associated with NCOR complex component TBL1XR1 and two components of the gene-silencing Polycomb group (PcG) multiprotein PRC1-like complex, RNF2 and PHC2 (Table 1).[63,68–70] These associations were only detected in untreated GBM cells, supporting the loss of the transcriptional repression activity of BCL6 in response to IR-Ac. However, in untreated glioblastoma cells, BCL6 was also commonly associated with Notch transcriptional coactivator MAML2 and NFκB transcriptional coactivator PIR, as well as transcriptional coregulator NCOA5 (Table 1).[67,71–77]

Therefore, while BCL6 associated with transcriptional corepressors in untreated glioblastoma cells, it also associated with transcriptional coactivators, suggesting potential for broad transcriptional activity. In response to IR-Ac, BCL6 ceased association with transcriptional coregulators in general, suggesting that BCL6 loses its transcriptional regulatory activity in response to IR-Ac treatment of glioblastoma cells.

Number of peptide spectral matches (PSMs) annotated to BCL6 and NCOR2 identified in each RIME replicate for each cell line and treatment.

### 2.6 Verification of BCL6 association with corepressor NCOR2 and subsequent IR-Ac-induced loss

The association of BCL6 with known corepressor NCOR2 in the untreated glioblastoma cell lines was verified using proximity ligation assays (PLAs). NFκB subunits p50 and p65 were used as a positive control. The co-localisation of p50 and p65 in LN18 cells was clearly differentiable from non-specific antibody controls (Fig. 5A). Similarly, the co-localisation of BCL6 and NCOR2 in all three glioblastoma cell lines was distinct (p ≤ 0.05) from the corresponding negative controls (Fig. 5B). This verified the association of BCL6 with NCOR2 in untreated glioblastoma cells.

**Figure 5:**
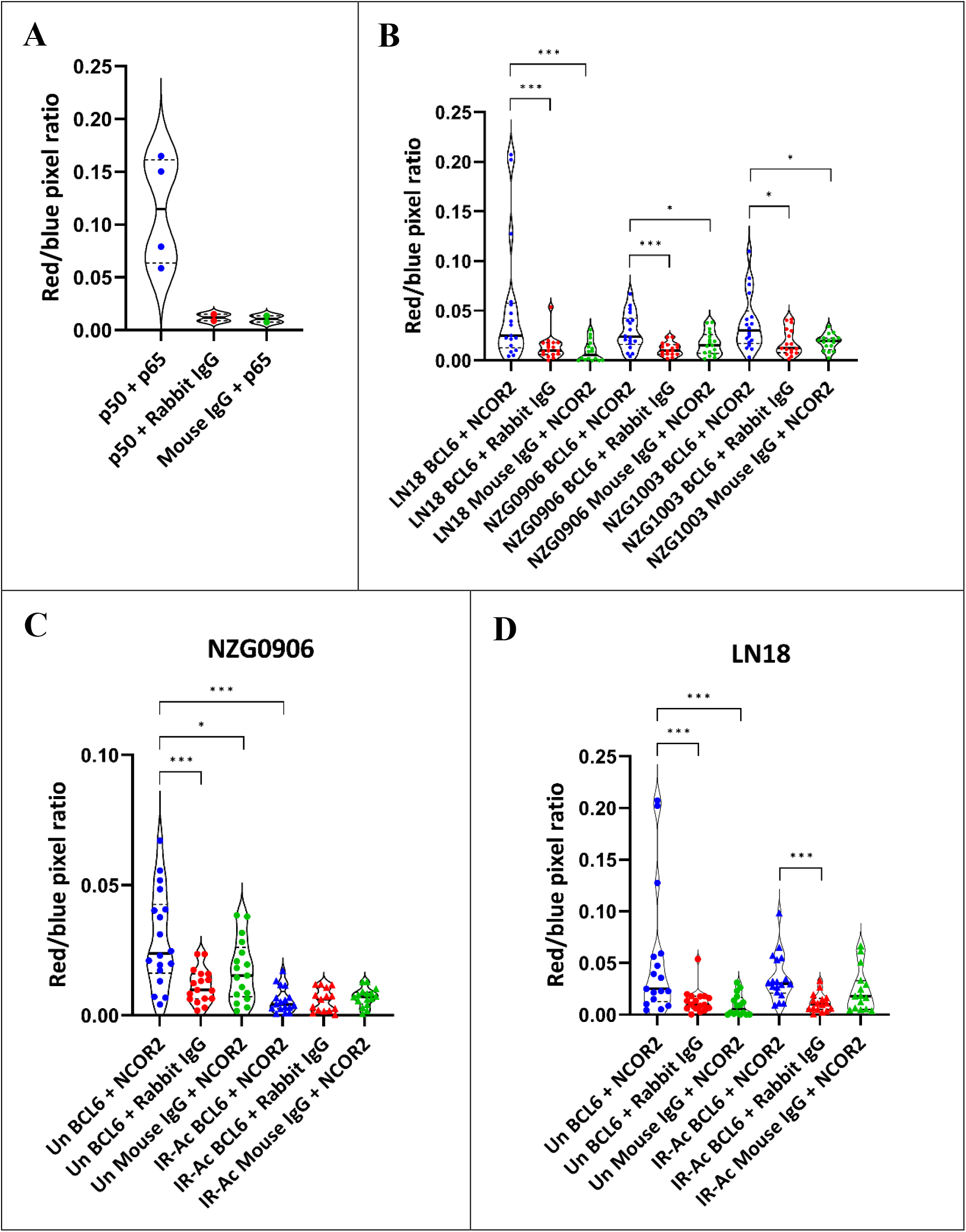
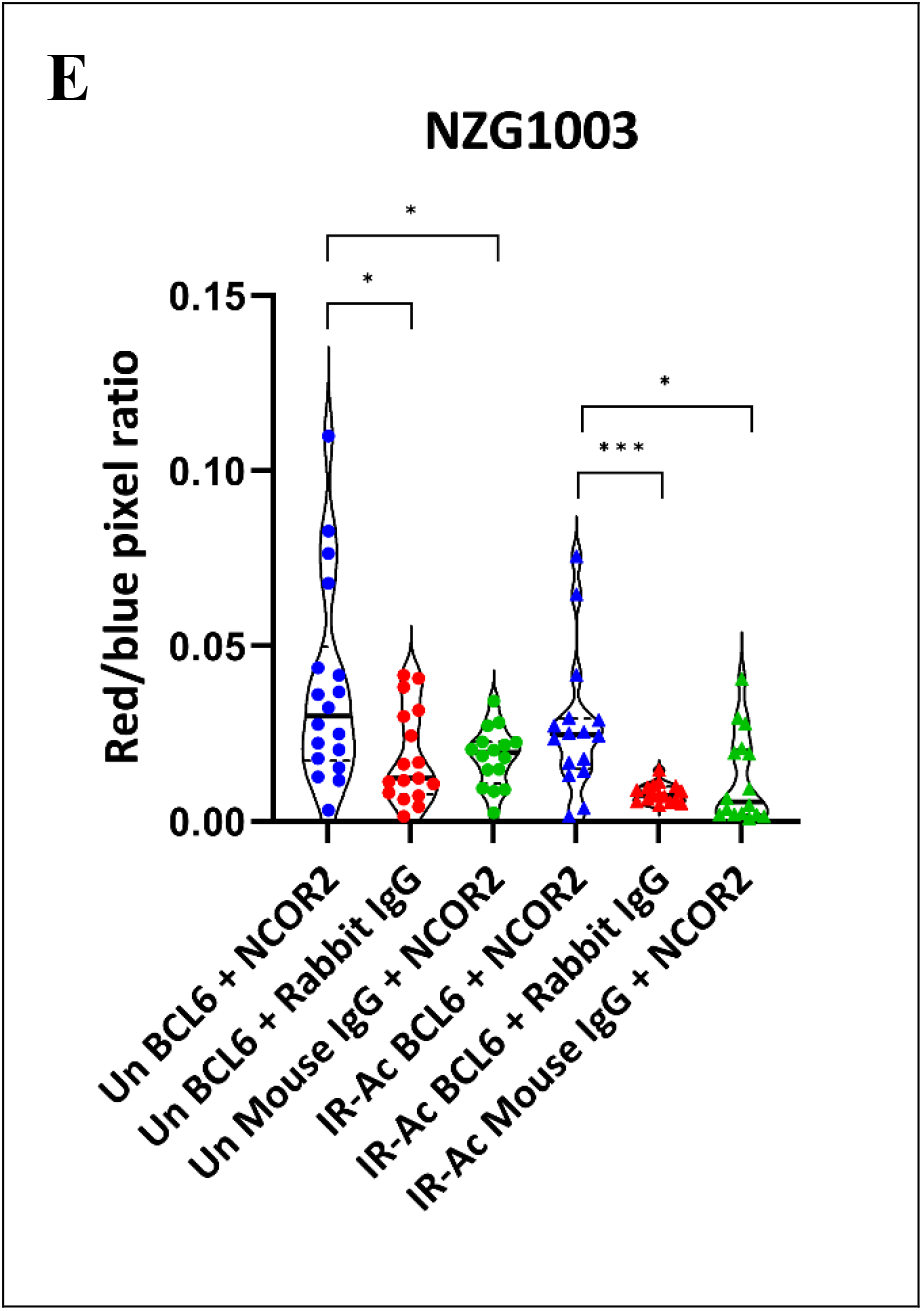
Verification of IR-Ac-induced loss of BCL6 association with NCOR2. **A)** Violin plots comparing PLA co-localisation signal (ratio of red to blue pixels) in NFκB (p50 + p65) positive control compared to negative control samples (p50 + non-specific rabbit IgG and p65 + non-specific mouse IgG) in LN18 cells. Bold line = median, dashed lines = quartiles. **B)** Violin plots comparing co-localisation signal (ratio of red to blue pixels) in BCL6 + NCOR2 proximity ligation assays compared to negative control samples (BCL6 + non-specific rabbit IgG and NCOR2 + non-specific mouse IgG) in untreated LN18, NZG0906 and NZG1003 cells. **C-E)** Violin plots comparing co-localisation signal (ratio of red to blue pixels) in BCL6 + NCOR2 proximity ligation assays compared to negative control samples (BCL6 + non-specific rabbit IgG and NCOR2 + non-specific mouse IgG) in untreated and irradiated **C)** NZG0906 cells, **D)** LN18 cells and **E)** NZG1003 cells. *** denotes p < 0.001, * denotes p < 0.05. Bold line = median, dashed lines = quartiles.

Next, the effect of IR-Ac on the BCL6-NCOR2 association was determined. Consistent with the RIME data, IR-Ac treatment of NZG0906 cells resulted in the loss of the BCL6-NCOR2 association (Fig. 5C) (p < 0.001). However, PLAs were unable to detect any change in BCL6-NCOR2 association in response to IR-Ac in LN18 (Fig. 5D) or NZG1003 (Fig. 5E) cells. Perhaps explaining this inconsistency, although the difference was statistically significant, the BCL6-NCOR2 interaction signal was only marginally higher than the signal in the non-specific antibody controls. This issue has been previously discussed and may indicate that the BCL6-NCOR2 interaction is close to the threshold of detection for this assay.[78]

## 3. Discussion

BCL6 is emerging as a critical protein for the survival of cancer cells. Research showing the importance of BCL6 in the survival and therapy resistance of glioblastoma has identified BCL6 as a promising target to improve the dire prognosis of this disease (Fig. 6).[11–15] This study demonstrates that the role of BCL6 in glioblastoma is highly context-specific, with dramatic changes in activity apparent in response to different levels of IR therapy (Fig. 6). In response to low dose, long-term IR-Fr treatment of glioblastoma cells, BCL6 was required for suppression of the DNA damage response, reminiscent of the canonical role of BCL6 in GC B cells.[29–31] However, in response to IR-Ac, BCL6 was upregulated and was required for the induction of a network of stress response signalling proteins, including p53, AKT1, NFκB1, AMPK-γ1 and cell cycle checkpoint proteins. BCL6 represses p53 and NFκB signalling in GC B cells and lymphoma, so this indicated a dramatic reversal of BCL6 activity in the response of LN18 glioblastoma cells to IR-Ac.[29–31] This suggested that as well as its known importance in adaptation to long-term stress, BCL6 also has a role in responses to single dose, high levels of stress.[79] However, the role of BCL6 in these two contexts appeared to be very different.

**Figure 6:**
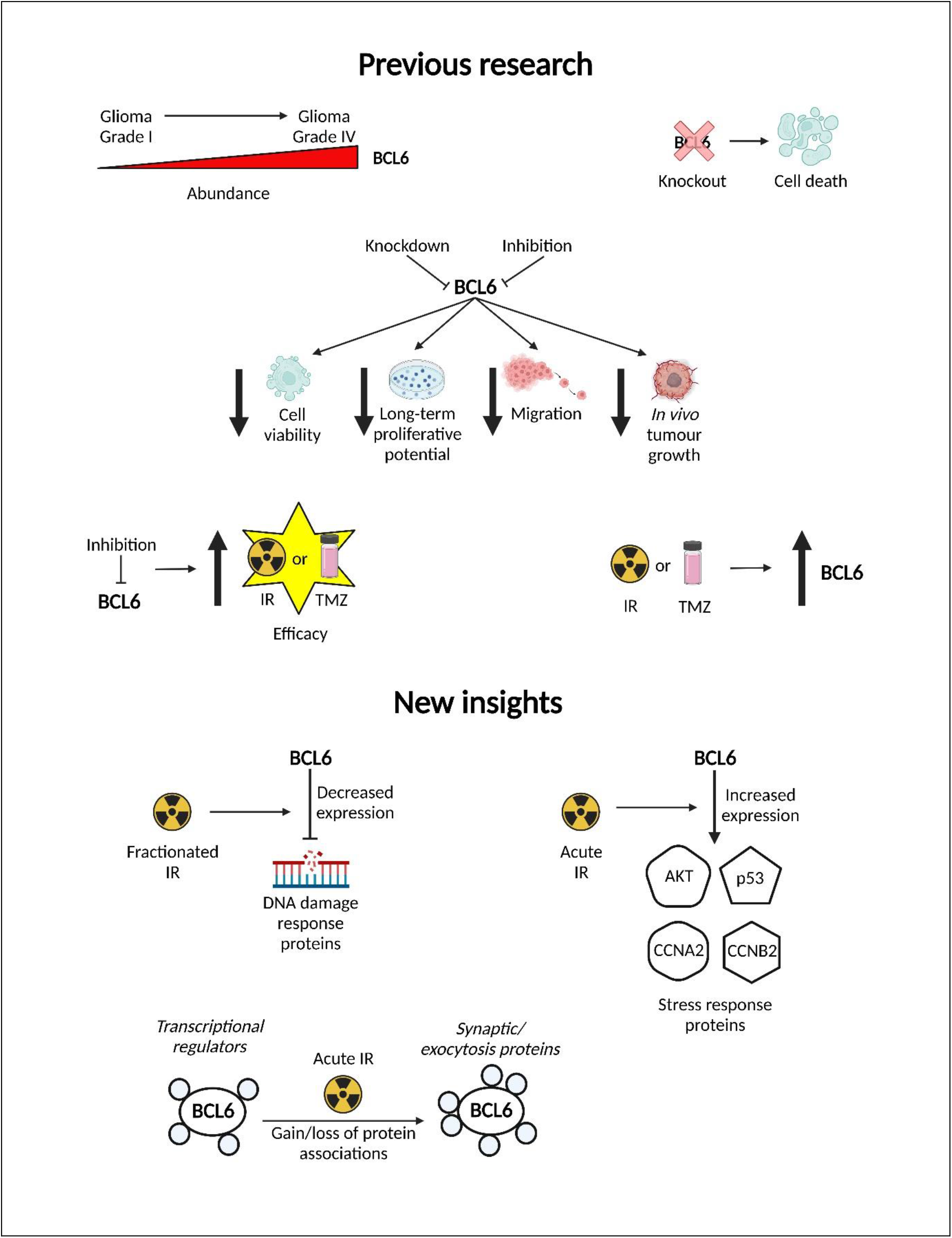
Summary of previous and new insights into BCL6 activity in glioblastoma. Previous research on BCL6 in glioblastoma is summarised in the top half of the figure, with new insights from the current study summarised in the bottom half. Created with BioRender.com.

The commercially available small molecule inhibitor FX1 was used to identify the BCL6-dependent component of the response to treatment. While commonly used as a BCL6 inhibitor and generally considered to be BCL6-selective, there is evidence that FX1 may affect other proteins, in particular CK2.[11,35,48,49,80–85] However, the data generated here from FX1-treated cells are largely consistent with the known role of BCL6. The apparent switch in BCL6 activity from repression of the DNA damage response to promotion of stress response signalling also fits with previous evidence that BCL6 may switch from a transcriptional repressor to a transcriptional activator in response to IR-Ac or doxorubicin treatment of glioblastoma cells.[11] However, it is possible that some of the proteins identified here are CK2-dependent rather than BCL6-dependent.

The canonical transcriptional activity of BCL6 is mediated and modified by the recruitment of corepressors.[17,19,24,86–88] RIME showed that in untreated glioblastoma cells, BCL6 associated with transcriptional regulators including known BCL6 corepressor NCOR2 and NCOR complex component TBL1XR1.[63] Additionally, BCL6 was associated with two Polycomb group proteins, RNF2 and PHC2, transcriptional activator of NOTCH signalling MAML2, coactivator of NFκB transcriptional activity PIR, and multifunctional transcriptional coregulator NCOA5.[67–71,76]

However, the association of BCL6 with transcriptional regulator proteins appeared to be lost after IR-Ac treatment. While identification of these protein associations fluctuated around the threshold of detection in LN18 cells, the IR-Ac-induced decrease in BCL6 association with NCOR2 was more distinct in NZG0906 cells, which expressed higher levels of BCL6. Similarly, PLA validation experiments supported the loss of the BCL6-NCOR2 association in response to IR-Ac in NZG0906 cells.

While the proteins associated with BCL6 changed dramatically in response to IR-Ac, there was no evidence that BCL6 gained association with transcriptional coactivators as previously proposed.[11] Instead, generalised loss of association with transcriptional regulators was evident. It is possible that in response to IR-Ac, BCL6 upregulates stress response signalling proteins indirectly rather than acting as a transcriptional activator itself. For example, the association of BCL6 with Notch and NFκB pathway coactivators MAML2 and PIR in untreated glioblastoma could indicate that BCL6 blocks transcriptional coactivation mediated by these proteins. The loss of BCL6 association with these coactivators after IR-Ac treatment may correspond to the release of this transcriptional blockade. The upregulation of stress response signalling was dependent on the BTB domain of BCL6, as it was blocked by FX1. It is possible that the interaction of proteins with the BTB domain is required for blocking interaction with MAML2 and PIR or for removal of BCL6 from the nucleus and association with other sub-cellular structures such as membrane vesicles. For example, the association of BCL6 with stress-responsive, pro-apoptotic kinase STK4 was increased after acute IR in LN18 and NZG0906 cells.[89]

Alternatively, BCL6 may interact with non-chromatin-associated proteins to promote upregulation of stress response signalling proteins. For example, after IR-Ac, BCL6 was associated with ubiquitin-binding protein UBXN1. UBXN1 suppresses NFκB signalling by sequestering cellular inhibitors of apoptosis proteins (cIAPs) to inhibit TNFα signalling and by sequestering CUL1 to prevent the degradation of NFKBIA.[90,91] If BCL6 association with UBXN1 disrupts the sequestering of these proteins, this could lead to activation of NFκB signalling. NFκB targets a wide variety of genes and release of NFκB inhibition could conceivably be the mechanism by which BCL6 upregulates stress response signalling.[92]

However, these speculations are inconsistent with the previously observed increase in the transcriptional activity of the BCL6 DNA binding motif in luciferase assays performed in treated glioblastoma cells.[11] Additionally, it must be noted that the decreased association of BCL6 with some proteins could have been affected by the lower overall number of protein detections in some IR-Ac-treated samples, which was due to the lower sensitivity of the mass spectrometer used to analyse these samples. This may have exacerbated the tendency of BCL6 and its associated proteins to fluctuate around the threshold of detection. Nevertheless, the enrichment of BCL6 afforded by RIME enabled new insights into the role of BCL6 in untreated and IR-Ac-treated glioblastoma. Further research is required to clarify the mechanism behind the change in BCL6 activity in response to IR-Ac.

The apparent localisation of BCL6 with synaptic proteins at the plasma membrane, including exocytic machinery, has intriguing implications. IR is known to increase extracellular vesicle (EV) release from glioblastoma cells.[93] In glioblastoma and other cancers, IR-induced EVs are taken up by surrounding cancer cells, where they induce bystander effects such as activation of DNA repair pathways, stress signalling and migration.[93–96] Therefore, it is possible that BCL6 is secreted in EVs in response to acute IR treatment to transmit stress responses important for cell survival to surrounding glioblastoma cells. Furthermore, exocytosis of BCL6 from irradiated glioblastoma cells could be a mechanism of the loss of BCL6-mediated transcriptional repression. It is also possible that BCL6 has another function at the plasma membrane, such as regulation of the exo- or endocytosis of signalling receptors. This could mediate the BCL6-dependent upregulation of AKT and NFκB signalling pathways in response to acute IR. Alternatively, as the whole proteomics analysis revealed that autophagy is induced by 48 hours after acute IR, it is also possible that BCL6 is caught up in autophagic vesicles with plasma membrane proteins rather than localising to the plasma membrane itself. This unexpected but striking finding will require further validation and investigation before any confident insights can be made.

BCL6 has been identified as an evolutionarily conserved stress response protein.[79] Canonically associated with repression of cellular stress responses in GC B cells, BCL6 is also important in the suppression of inflammatory phenotypes in T_FH_ cells, T_regs_ and macrophages.[29–31,97–100] Cancer cells have inherently stressful cellular environments due to their high rates of proliferation, mutational burdens and altered metabolism.[79] BCL6 expression is upregulated in multiple cancer types and its inhibition has been shown to increase the efficacy of stress-inducing cancer treatments.[11,49,79,80] This study demonstrates that the role of BCL6 in the stress response of cancer cells may be dynamic and may alter depending on the type or level of stress. BCL6 appeared to repress DNA damage repair in response to IR-Fr treatment of LN18 GBM cells. However, when BCL6 was upregulated by IR-Ac, it appeared to switch to promoting stress response signalling and pathways that promote DNA damage repair. This suggests that BCL6 may play a different role in the response to acute, high levels of stress than it does in adaptation to long-term stress. The suppression of the DNA damage response may be beneficial to cancer cells when it enables them to continue to proliferate rather than undergoing cell cycle arrest or apoptosis. However, if DNA damage gets too severe, this may no longer be beneficial. Instead, the upregulation of stress response signalling could allow these cells to repair the damage and survive. Hence, the ability to either suppress or change BCL6 function in response to high levels of stress could be an important component of the BCL6-mediated stress response pathway.

In summary, the role of BCL6 in the therapy response of glioblastoma cells is more complicated and context-specific than expected. Evidence is mounting that BCL6 is important for the survival of glioblastoma cells and for their resistance to treatment, although more research is needed to fully clarify the role of BCL6 in the survival response of glioblastoma cells to different treatments. However, BCL6 inhibitors in development hold great promise to improve the efficacy of the gold-standard treatment for people diagnosed with this aggressive malignant disease.

## 4. Methods

### 4.1 Cell lines

The human glioblastoma cell lines LN18, NZG0906 and NZG1003 were grown as adherent cultures in RPMI 1640 media supplemented with 5% foetal bovine serum. The LN18 cell line was sourced from the American Type Culture Collection (USA). The NZG0906 and NZG1003 cell lines were previously immortalised in the McConnell lab from tumour tissue.[60] The Raji human lymphoma cell line was a gift from Ian Morison, University of Otago, and was cultured in suspension in RPMI 1640 supplemented with 5% FBS and 1 mM pyruvate. All cell lines were grown under conditions of normoxia, with 5% CO2 and at 37 °C.

### 4.2 Western blot

LN18 cells were lifted and frozen at –80 °C. The cells were thawed, lysed in 8 M urea and quantified using the Pierce™ Rapid Gold BCA Protein Assay Kit (Thermo Fisher Scientific, USA). Samples were denatured in Laemmli buffer at 95 °C and run at 170 V on a 10% acrylamide gel with a 4% stacking gel. Transfer to a PVDF membrane was achieved at 300 mA for 2 hours. The membrane was blocked in 5% milk powder in PBS at room temperature for 1 hour, followed by incubation with a 1:1000 dilution of the anti-BCL6 D-8 antibody (Santa Cruz, USA) in the same solution overnight at 4 °C. 0.1% Triton X-100 in PBS was used to wash the membrane. A 1:7000 dilution of HRP goat anti-mouse IgG antibody (BioLegend, USA) was applied for 1 hour at room temperature in 0.1% Triton X-100 in PBS. After final washes, the membrane was imaged using Western Lightning Ultra Chemiluminescent Substrate (Perkin-Elmer, USA) and an Amersham Imager 600 (GE Health Life Sciences, USA) The membrane was stripped and re-stained using an anti-β-actin AC-15 antibody (Sigma Aldrich, USA). Western blot bands were quantified relative to the β-actin loading control using Fiji (ImageJ).[101]

### 4.3 Protein preparation for whole proteomics

In biological triplicate, LN18 cells were treated with either 2 Gy irradiation every day for 5 doses (IR-Fr) or with one dose of 10 Gy irradiation (IR-Ac). Additionally, the LN18 cells were treated with 10 μM FX1 or the equivalent volume of DMSO on the day of 10 Gy irradiation and on the first, third and fifth days of 2 Gy irradiation. IR-Fr-treated cells were passaged as required during the treatment regime. Untreated cells were treated with the same volume of DMSO only. The cells were harvested and frozen 24 hours after the final treatment.

Cell pellets were thawed and lysed in 8 M urea. After acetone precipitation, the dried pellet was resuspended in 8 M urea and quantified using the Pierce™ Rapid Gold BCA Protein Assay Kit (Thermo Fisher Scientific, USA). A 2 hour incubation with 10 mM DTT at 56 °C was followed by 40 mM iodoacetamide for 45 minutes in the dark at room temperature. After dilution of the samples to 2 M urea, Pierce™ trypsin protease, MS grade (Thermo Fisher Scientific, USA) was added in a weight ratio of 1:100 overnight at 37 °C.

### 4.4 RIME

Each RIME sample was derived from 10 confluent 15 cm plates of LN18, NZG0906 or NZG1003 cells. For 10 Gy irradiation treatment, cells were initially plated into 20 10 cm plates before being lifted and transferred into 15 cm plates 24 hours after treatment. Cells were left for another 24 hours before being processed for RIME (48 hours after irradiation). The untreated cells were also lifted and replated 24 hours before processing.

The rapid immunoprecipitation mass spectrometry of endogenous proteins (RIME) protocol published by Mohammed et al. (2016) was performed with a few modifications.[39] All procedures, except sonication, were performed in a Laminar Flow hood.

For each RIME sample, 1 mg Invitrogen™ Protein G Dynabeads™ (Thermo Fisher Scientific, USA) was added to ten low protein binding microcentrifuge tubes. The beads were washed and then resuspended in 40 µL RIPA buffer (50 mM HEPES (pH 7.6), 1 mM EDTA, 0.7% (w/v) sodium deoxycholate, 1% (v/v) NP-40, 0.5 M LiCl). 5 µg N-3 anti-BCL6 antibody (Santa Cruz, USA) was added to five of the tubes and 5 µg non-specific Invitrogen™ normal rabbit IgG antibody (Thermo Fisher Scientific, USA) was added to the other five tubes. The tubes were rotated at 4 °C while the cells were prepared.

Cells were fixed with 1% (v/v) formaldehyde for 8 minutes, then quenched with 0.125 M glycine for 10 minutes. After washes in cold PBS, cells were scraped into 10 mL PBS per plate and pelleted at 2000 *g* for 10 minutes at 4 °C. 10 mL swelling buffer (50 mM HEPES-KOH (pH 7.5), 140 mM NaCl, 10% glycerol, 1 mM EDTA, 0.5% (v/v) IGEPAL, 0.25% (v/v) Triton X-100) was added to the pooled RIME sample with fresh Halt™ Protease Inhibitor Cocktail (Thermo Fisher Scientific, USA). 1 mL aliquots of the sample were rotated at 4 °C for 10 minutes before the nuclei were pelleted at 1700 *g* for 5 minutes at 4 °C. The pellets were resuspended in 1 mL wash buffer (10 mM Tris-Cl (pH 8.0), 200 mM NaCl, 1 mM EDTA, 0.5 mM EGTA) with fresh protease inhibitor. The samples were rotated and pelleted as previously.

Pellets were resuspended in 99.25 µL micrococcal nuclease reaction buffer (1:10 micrococcal nuclease 10 x buffer (NEB), 1:100 10 mg/mL BSA in distilled water) with 0.75 µL (150 U) micrococcal nuclease enzyme solution (1:10 micrococcal nuclease (NEB, USA) in micrococcal nuclease reaction buffer). The reactions were incubated at 37 °C for 15 minutes, with vortexing every two minutes, before addition of 10 µL 0.5 M EDTA and placement onto ice. Nuclei were pelleted at 16,000 *g* for 1 minute at 4 °C and resuspended in 200 µL shearing buffer (50 mM Tris-Cl (pH 8.1), 0.1% (w/v) SDS, 10 mM EDTA (pH 8.0)) with fresh protease inhibitor and incubated on ice for 10 minutes. Sonication with 3 x 20 second pulses with 30 seconds rest in between further fragmented the DNA. The lysates were clarified at 9400 *g* for 10 minutes at 4 °C and then pooled and diluted 1:1 in dilution buffer (50 mM Tris-Cl (pH 8.0), 300 mM NaCl, 2% (v/v) IGEPAL, 1% (w/v) sodium deoxycholate (Na-DOC), 0.1% (v/v) sodium dodecyl sulfate (SDS)). The sample was distributed evenly between the 10 prepared antibody/Dynabead™ tubes which were rotated overnight at 4 °C.

The supernatant was discarded and beads were washed in 1 mL RIPA buffer and rotated at 4 °C for 5 minutes 10 times. After a final two washes in 1 mL fresh 100 mM ammonium hydrogen carbonate, the beads were resuspended in 10 µL 100 mM ammonium hydrogen carbonate containing 100 ng Pierce™ trypsin protease, MS grade (Thermo Fisher Scientific, USA) and incubated at 37 °C overnight. A further 10 µL 100 mM ammonium hydrogen carbonate containing 100 ng trypsin was added for a further 4 hour incubation at 37 °C the next day. The beads were pelleted on a magnet and the supernatant was retained. The samples treated with the same antibody (BCL6 or IgG) were pooled.

### 4.5 Peptide desalting

Formic acid was added to all peptide samples to 0.1%, before desalting using BondElut OMIX 100 µL C18 tips. The desalted peptides were dried and resuspended in HPLC-grade water for quantification using the Pierce™ Quantitative Fluorometric Peptide Assay (Thermo Fisher Scientific, USA).

### 4.6 Mass spectrometry analysis

Due to instrument availability, some samples were run on the Orbitrap Fusion™ Lumos™ Tribrid™ Mass Spectrometer (Thermo Fisher Scientific, USA) at Victoria University of Wellington, New Zealand, while others were dried and sent to the Bio21 Institute at the University of Melbourne, Australia for analysis.

Samples run on the Lumos™ were resuspended at 100 ng/µL in 0.1% formic acid in HPLC-grade water. The settings of the autosampler and mass spectrometer were defined using Xcalibur™ 4.2 software (version 2.1.0). For the whole proteomics samples, 200 ng of each peptide sample was loaded by the Dionex UltiMate 3000 RS Autosampler for separation by liquid chromatography, in technical duplicate. For the RIME samples, 100 ng of each peptide sample was loaded. The peptides were loaded onto an Acclaim™ PepMap™ 100 C18 trap column (5 µm, 0.3 x 5 mm) with 0.05% formic acid in 2% acetonitrile at a loading pump flow rate of 8 µL/minute. The peptides were then separated on an Acclaim™ PepMap™ 100 C18 analytical column (2µm, 100 A, 75 µm x 15 cm).

The nanocolumn (NC) pumped Buffers A (0.1% formic acid in HPLC-grade water) and B (0.1% formic acid in 80% acetonitrile) at 0.3 µL/minute. For the whole proteomics samples, the flow gradient was (i) 0-5 minutes at 3% B, (ii) 5-70 minutes from 3-30% B, (iii) 70-80 minutes from 30-50% B, (iv) 82-83 minutes from 50-95% B, (v) 83-88 minutes at 95% B, (vi) 88-90 minutes from 95-3% B, (vii) 90-99 minutes at 3% B. For the RIME samples, the flow gradient was (i) 0-5 minutes at 3% B, (ii) 5-10 minutes from 3-10% B, (iii) 10-45 minutes from 10-25% B, (iv) 45-50 minutes from 25-50% B, (v) 50-51 minutes from 50-95% B, (vi) 51-56 minutes at 95% B, (vii) 56-57 minutes from 95-3% B, (viii) 57-70 minutes at 3% B. The column was washed between every two samples with the following gradient: 0-5 minutes at 3% B, 5-6 minutes from 3-95% B, 6-9 minutes at 95% B, 9-10 minutes 95-3% B. This gradient was repeated three times with a final 14 minutes at 3% B.

The peptides eluted from the column were injected into the mass spectrometer using nanospray ionisation. The ion transfer tube (25 µm) was set to 275 °C and the voltage was set to 1800 V. The MS1 scans were obtained in positive mode using quadrupole isolation with a scan range of 375-1500 m/z and with detection at a resolution of 120,000 in the Orbitrap. The maximum injection time was 50 ms and the normalised automatic gain control (AGC) target was 175%. From each MS1 scan, the 20 highest intensity ions were selected for MS2 scans on ions in an isolation window of 1.6 m/z, with charge 2-7 and intensity above 5.0E3. These ions were selected using the quadrupole, with a dynamic exclusion duration of 60 seconds after a single detection and a mass tolerance of ±10 ppm. Precursor ions were fragmented using higher energy C-trap dissociation (HCD) with an HCD collision energy of 30%. Ions were detected in the ion trap with a dynamic maximum injection time and a normalised AGC target of 50%.

The following details were provided by the Bio21 Institute upon request. The LC system was equipped with an Acclaim Pepmap nano-trap column (Dinoex-C18, 100 Å, 75 μm x 2 cm) and an Acclaim Pepmap RSLC analytical column (Dinoex-C18, 100 Å, 75 μm x 50 cm). The tryptic peptides were injected to the enrichment column at an isocratic flow of 5 μL/min of 2% v/v CH_3_CN containing 0.05% v/v TFA for 6 min applied before the enrichment column was switched inline with the analytical column. The eluents were 5% DMSO in 0.1% v/v formic acid (solvent A) and 5% DMSO in 100% v/v CH_3_CN and 0.1% v/v formic acid (solvent B). The flow gradient was (i) 0-6min at 3% B, (ii) 6-40min, 3-25% B (iii) 40-48min 25-45% B (iv) 48-50min, 45-80% B (v) 50-53in, 80-80% B (vi) 53-54min, 80-2% and equilibrated at 2% B for 10 minutes before the next sample injection. The Q Exactive Plus™ mass spectrometer was operated in the data-dependent mode, whereby full MS1 spectra were acquired in positive mode, 70 000 resolution, AGC target of 3e_6_ and maximum IT time of 50ms. Fifteen of the most intense peptide ions with charge states ≥ 2 and intensity threshold of 4e_4_ were isolated for MSMS. The isolation window was set at 1.2m/z and precursors fragmented using normalized collision energy of 30, 17 500 resolution, AGC target of 5e_4_ and maximum IT time of 50ms. Dynamic exclusion was set to be 30sec.

### 4.7 Protein identification from mass spectrometry data

Raw mass spectra data files were uploaded to Proteome Discoverer 2.4 and searched against the Swiss-Prot reviewed UniProt human protein database.[102] MS1 precursors were filtered by mass (350-5000 Da), minimum peak count (1), length (6-144 amino acids), maximum missed cleavage sites (2), precursor mass tolerance (10 ppm), fragment mass tolerance (0.5 Da) and maximum RT shift (10 minutes).

Carbamidomethyl modifications (static) and oxidation of methionine and deamidation of asparagine and glutamine (dynamic) were reported for whole proteomics and RIME samples respectively if site probability was ≥75%. Protein groups were formed using the strict parsimony principle and master proteins were assigned. A decoy search was performed by the Percolator node with concatenated target/decoy selection.

### 4.8 Protein quantification from whole proteome mass spectrometry data

Peak intensity-based quantification used both unique and razor peptides followed by normalisation to total peptide amount and scaling of abundances so the average of all samples was 100. Protein abundances were calculated from summed peptide abundances and were compared with pairwise ratios by taking the median of all possible pairwise peptide ratios between the replicates. Fold-changes were capped at 100 and statistical significance of quantification ratios was assessed by background-based t-tests. The grouped biological replicates for each treatment were compared to other treatments in non-nested ratio analyses. Low confidence proteins (exp. q value ≥ 0.05) were excluded from downstream analysis.

### 4.9 Protein quantification from RIME mass spectrometry data

Quantification was performed as above (section 4.8) except for RT tolerance (0.0001 minutes) and mass tolerance (1E-5 ppm), which were set low to effectively prevent the match between runs (MBR) function which otherwise led to aberrant identification of BCL6 in the IgG samples. Proteins included in downstream analysis were high confidence (exp. q value ≤ 0.01), upregulated ≥ 2-fold (p ≤ 0.05) and not found at high abundance in any IgG samples.

Protein abundances were normalised to the abundance of BCL6, as defined by a BCL6 amino acid sequence FASTA file. Protein abundance ratios were calculated directly from the grouped protein abundances.

### 4.10 Functional enrichment analysis

UniProt protein accession numbers were entered into g:Profiler version *e104_eg51_p15_2719230*.[102,103] Any protein accession numbers not recognised by g:Profiler were converted into Ensembl gene (ENSG) IDs if available.[104] Obsolete Uniprot accession numbers were replaced with the new accession number if available. Any remaining proteins were excluded from the g:Profiler analysis. Statistical domain scope was set to ‘only annotated genes’, significance threshold to ‘g:SCS threshold’ and user threshold to 0.05. Data sources selected were the Gene Ontology categories GO molecular function (GO:MF), GO biological process (GO:BP) and GO cellular component (GO:MF).

Network analysis was performed using STRING version 11.5, utilising interaction sources textmining, experiments, databases, co-expression, neighbourhood and cooccurrence.[46] Edge thickness indicated confidence and the minimum required confidence was high (0.7).

### 4.11 Proximity ligation assays

Cells were plated into chamber slide wells 24 hours before staining and 24 hours after 10 Gy irradiation treatment when applicable. Cells were washed with PBS and fixed in 4% PFA for 15 minutes at room temperature. After further PBS washes, the cells were permeabilised in PBS, 0.1% Triton X-100 for 15 minutes on ice. Cells were stained as described in the Duolink® PLA Fluorescence Protocol (Merck, USA).[105,106] Antibodies were added in Antibody Diluent: 1:50 (4 µg/mL) D-8 anti-BCL6 (Santa Cruz, USA); 1:150 (6.7 µg/mL) anti-NCOR/SMRT (Abcam, UK); 1:500 (1 µg/mL) 4D1 anti-p50 (BioLegend, USA); 1:500 (0.2 µg/mL) anti-p65 (Sigma Aldrich, USA); 4 µg/mL eBioScience™ mouse IgG (Thermo Fisher Scientific, USA); 6.7 µg/mL Invitrogen™ normal rabbit IgG (Thermo Fisher Scientific). For the amplification step, incubation at 37 °C was kept to 100 minutes for the NFκB positive control, as per the Duolink® protocol, but was extended to 3.5 hours for the BCL6 and NCOR2 experiments.

Images were acquired using an Olympus Laser Scanning Confocal Microscope (LSCM) FV3000 in an inverted microscope frame IX83 (Malaghan Institute of Medical Research). Images were taken as 10 z-stacks through the whole depth of the cells as assessed by DAPI staining of the nuclei. Acquisition settings were the same for all samples which were directly compared. Other settings were as follows: 137 μm confocal aperture; 1024×1024 pixel images; 1% laser power; 1x gain; DAPI excitation at 405 nm, 430 V, 5% offset; PLA fluorophore (Texas Red®) excitation at 561 nm, 350 V, 5% offset.

### 4.12 Image analysis

Images were converted to composite z-projections using Fiji (ImageJ).[101] The relative brightness of each channel was adjusted (consistently between images) using CellProfiler.[107] Red and blue pixels were counted in Fiji (ImageJ) using the Color Pixel Counter plugin, with minimum intensity thresholds set to 50.[108] Statistical comparisons were made with SPSS using a linear mixed model with antibody pairs and treatment as fixed effects, replicates as a random effect and images within replicates as repeated measurements with compound symmetry. Pairwise comparisons between antibody pairs and between treatments were made using sequential Bonferroni multiple comparisons adjustment.

## Data accessibility

The commonly identified BCL6-associated proteins were submitted to the IMEx (http://www.imexconsortium.org) consortium through IntAct [X] and assigned the identifier IM-29565.[109]

Mass spectra raw files and peak lists and Proteome Discover result files are available on the MassIVE database version 1.3.16 (ftp://MSV000090274@massive.ucsd.edu).[110]

## Acknowledgements

The guidance of Alfonso Schmidt (Malaghan Institute of Medical Research, New Zealand) regarding confocal microscopy and the advice of Soleilmane Omarjee (Cancer Research UK – Cambridge Institute) and Hamish McMillan (Otago University, New Zealand) regarding the RIME protocol are acknowledged and appreciated. The Malaghan Institute is also acknowledged for providing access to their irradiator and confocal microscope.

## FIG

